# Rapid *in vitro* method to assemble and transfer DNA fragments into the JCVI-syn3B minimal synthetic bacterial genome through Cre/*loxP* system

**DOI:** 10.1101/2024.08.01.606163

**Authors:** Atsuko Uenoyama, Hana Kiyama, Mone Mimura, Makoto Miyata

**Author notes:** **Corresponding author:** Makoto Miyata, Graduate School of Science, Osaka Metropolitan University, Osaka, 558-8585, Japan.

## Abstract

JCVI-syn3B (syn3B), a minimal synthetic bacterium that only possesses essential genes, facilitates the examination of heterogeneous gene functions in minimal life. Conventionally, *Escherichia coli* is used to construct DNA fragments for gene transfer into the syn3B genome through Cre/*loxP* system. However, the construction process is challenging and time-consuming due to various issues, including the inhibition of *E. coli* growth and unexpected recombination, especially with AT-rich DNA sequences such as those found in *Mycoplasma* genes. Therefore, in this study, we aimed to develop a new transformation method to overcome these issues. We assembled the vector and target DNA fragments using an in vitro homologous recombination system and subsequently transferred the products into the syn3B genome. We obtained approximately 10^3^∼10^4^ recombinant colonies per milliliter of the original culture in eight days, which is four days shorter than the conventional period, without any recombination issues, even for AT-rich DNA. This method may be applicable to other gene manipulation systems based on Cre/*loxP* system.

**Significance:** A rapid and trouble free method was developed to transfer genes to the genome of minimal synthetic bacterium JCVI-Syn3B through Cre/*loxP* system. This method can be applied to Cre/*loxP*-based gene manipulation system in various research fields.

## Introduction

JCVI-syn3B (syn3B) is a “synthetic” bacterium based on a small bacterium, *Mycoplasma mycoides*, which was established to answer the question what is the minimal life? The necessity of individual *M. mycoides* genes were judged by bioinformatics and experiments. The syn3B genome was designed by a computer, synthesized chemically, assembled in *E. coli* and yeast, and then transplanted to a related species, *Mycoplasma capricolum* [1]. The syn3B genome contains a locus of *loxP* site to which external genes can be efficiently transferred via the Cre recombinase activity (Fig. 1) [2-4]. Therefore, syn3B is a useful platform for both basic and applied researches [3,5-8]. Our group is discussing the origin of cell motility, based on syn3B experiments [5,9]. In the transformation of syn3B by external genes requires a plasmid containing the target DNA, Cre recombinase, *loxP* sites, and puromycin resistance gene [2]. We construct these plasmids using *Escherichia coli* (Fig. 1) as described below. (a, day 1) Polymerase chain reaction (PCR)-amplified vector and target DNA fragment were assembled via homologous recombination-based methods using the NEBuilder® and In-Fusion® systems. The constructs were then used to transform *E. coli* cells. (b, day 2) Transformed colonies were inoculated into a liquid medium. (c, days 3–5) Plasmids were isolated and sequenced. (d, days 6–9) Syn3B was transformed with the plasmid. (e, days 10–12) Colonies were cultured in a liquid medium and target gene transfer was confirmed via direct PCR. This method exhibited some drawbacks, including poor *E. coli* growth and unexpected recombination. Moreover, we suffered from unexpected gene recombination in transfer of some *Mycoplasma* and related species, possibly due to their high AT content (67–76%) [10,11]. Therefore, in this study, we aimed to explore four transfer methods without *E. coli* to develop a new *E. coli*-free approach that is four days faster than the conventional method for gene transfer.

**Fig. 1.**
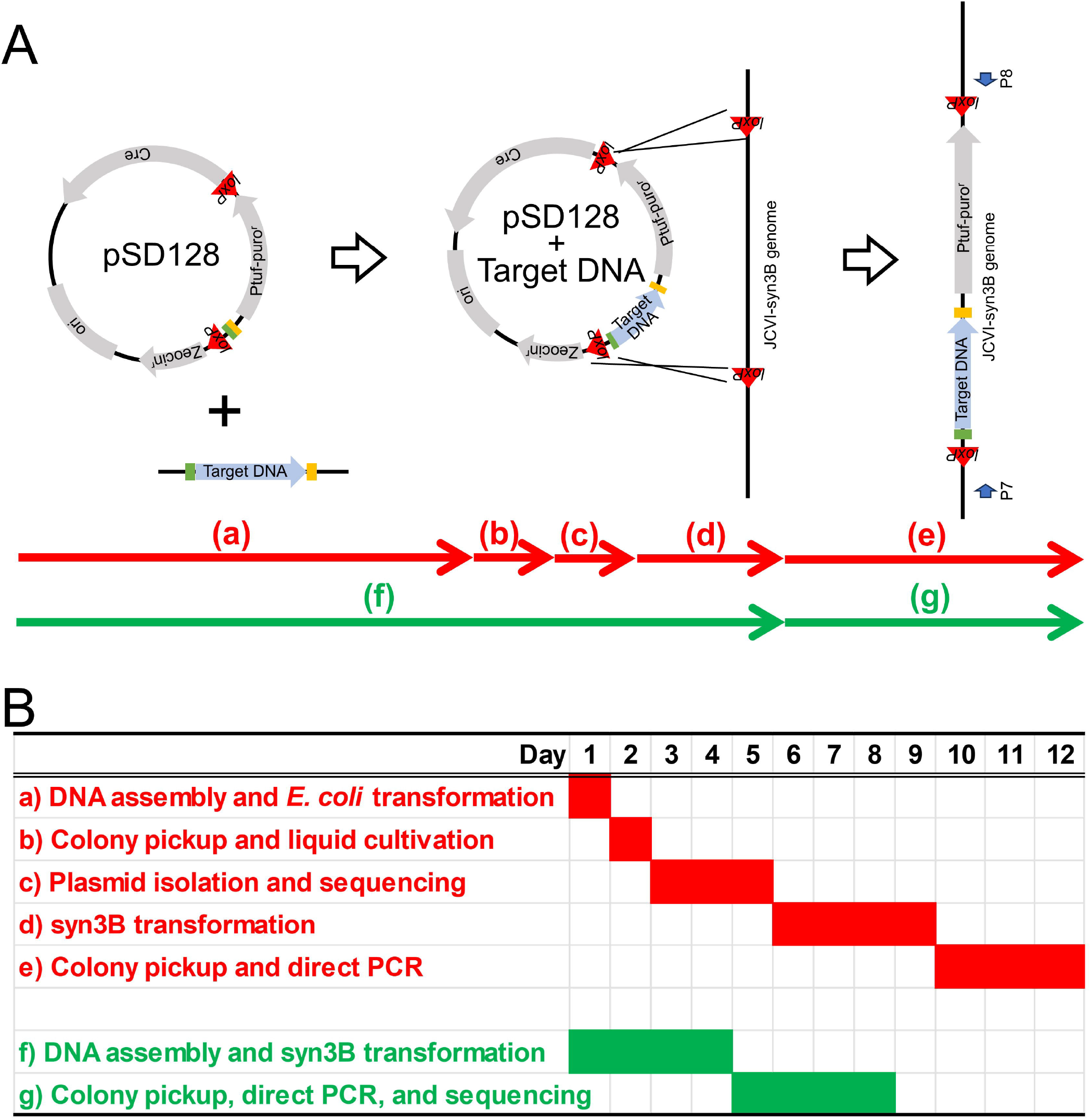
DNA transfer into the JCVI-syn3B (syn3B) genome. (A) Schematic illustration. Plasmid pSD128 used for gene transfer contains a Cre/*loxP*-related region and puromycin resistance gene. Target DNA is transferred into the syn3B genome via the Cre recombinase activity. Blue arrows indicate the primers used for direct polymerase chain reaction (PCR). The conventional (red) and new (green) processes are shown as five (a–e) and two (f–g) steps, respectively. (B) Gantt chart for the conventional (red) and new (green) processes.

## Materials and Methods

### Bacterial strains and culture conditions

JCVI-syn3B (GenBank, CP069345.1) and *E. coli* (DH5α) were cultured in the SP4 [1,2] and Luria–Bertani media at 37°C, respectively.

### Preparation of DNA fragments

*Mycoplasma mobile* genome was isolated as previously described [12]. Vector and focused *M. mobile* DNA fragments were amplified from pSD128 DNA [3] and *M. mobile* genomic DNA using primers P1–P2 and P3–P4 (Table 1), respectively. The DNA fragments were assembled using the NEBuilder® HiFi DNA Assembly Master Mix (New England Biolabs, Inc., MA, USA) and In-Fusion HD® Cloning Kit (Takara Bio Inc., Kusatsu, Japan). Then, the assembled product was amplified via PCR using primers P5 and P6 and circularized via self-assembly with NEBuilder®.

**Table 1.**
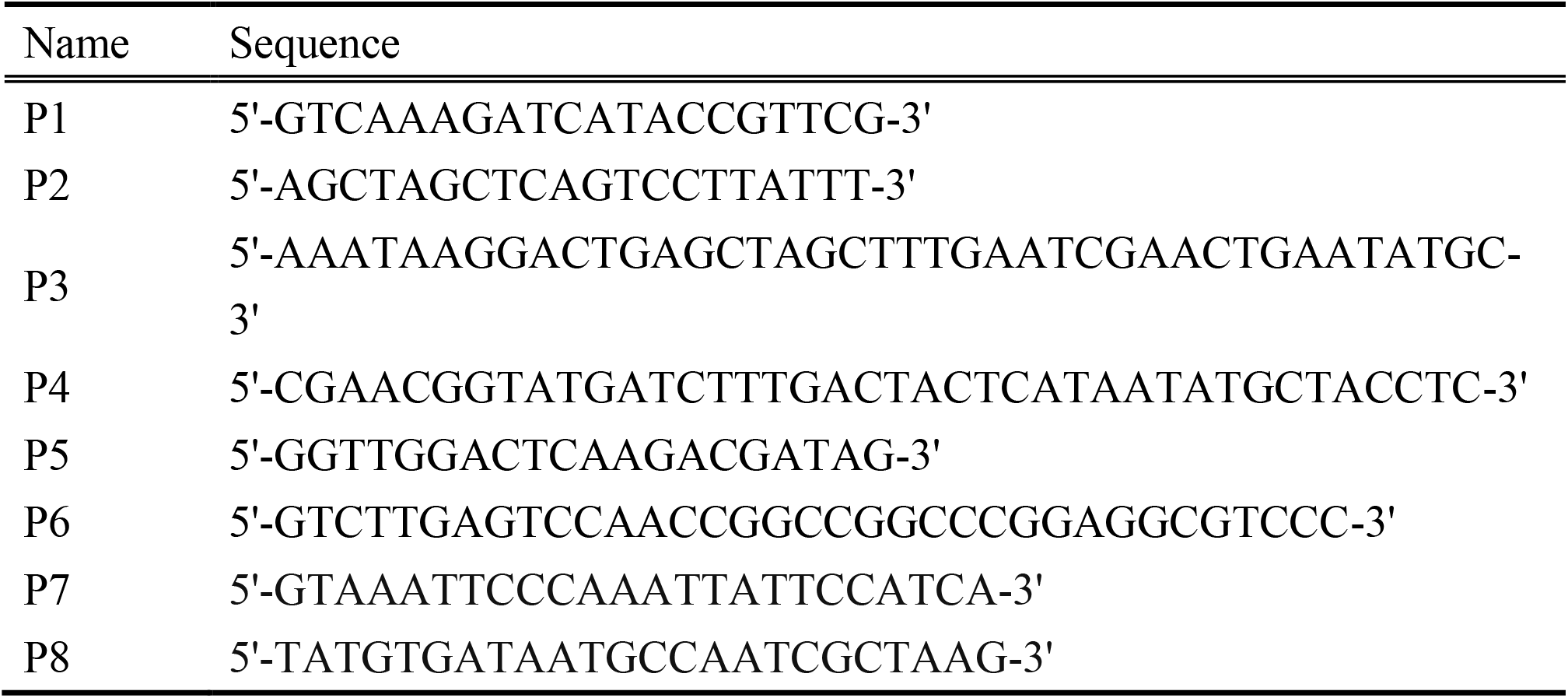
Primers used in this study.

### Transformation and direct PCR

Transformation of JCVI-syn3B was performed as previously described [5]. In short, competent cells were incubated with respective DNA for transfer for 15 min on ice, treated with 70% PEG_6000_, and then incubated in SP4 medium at 37°C for 3 h. Cells were then plated on a SP4 (containing horse serum instead of FBS) plate supplemented by 3 µg/mL puromycin. After incubation at 37°C for four days, the number of transformed colonies was counted. The transformation efficiency is shown as values relative to those of 100 ng pSD128 plasmid. This is because the efficiency largely depended on serum. For further studies, the transformed colonies were picked and inoculated into 100 µL of SP4 medium containing puromycin (3 µg/mL) for 18–24 h to make stocks, and confirmed via PCR using primers P7 and P8.

## Results and Discussion

### Rationale of the transformation protocol

To skip DNA manipulation in *E. coli*, we assembled the DNA fragments using an in vitro homologous recombination system and amplified them using PCR to obtain large amount of uniform DNA fragments, beneficial for transformation. Four types of DNAs were tried as shown in Fig. 2. As our target DNA, we used a 1,916-bp fragment containing the 1,536-bp gene encoding phosphoglycerate kinase from *M. mobile* (MMOB4530) and its flanking region (at 565,247–567,161 nt on the genome) [13-17]. **Transformation using assembled and amplified linear DNA**

**Fig. 2.**
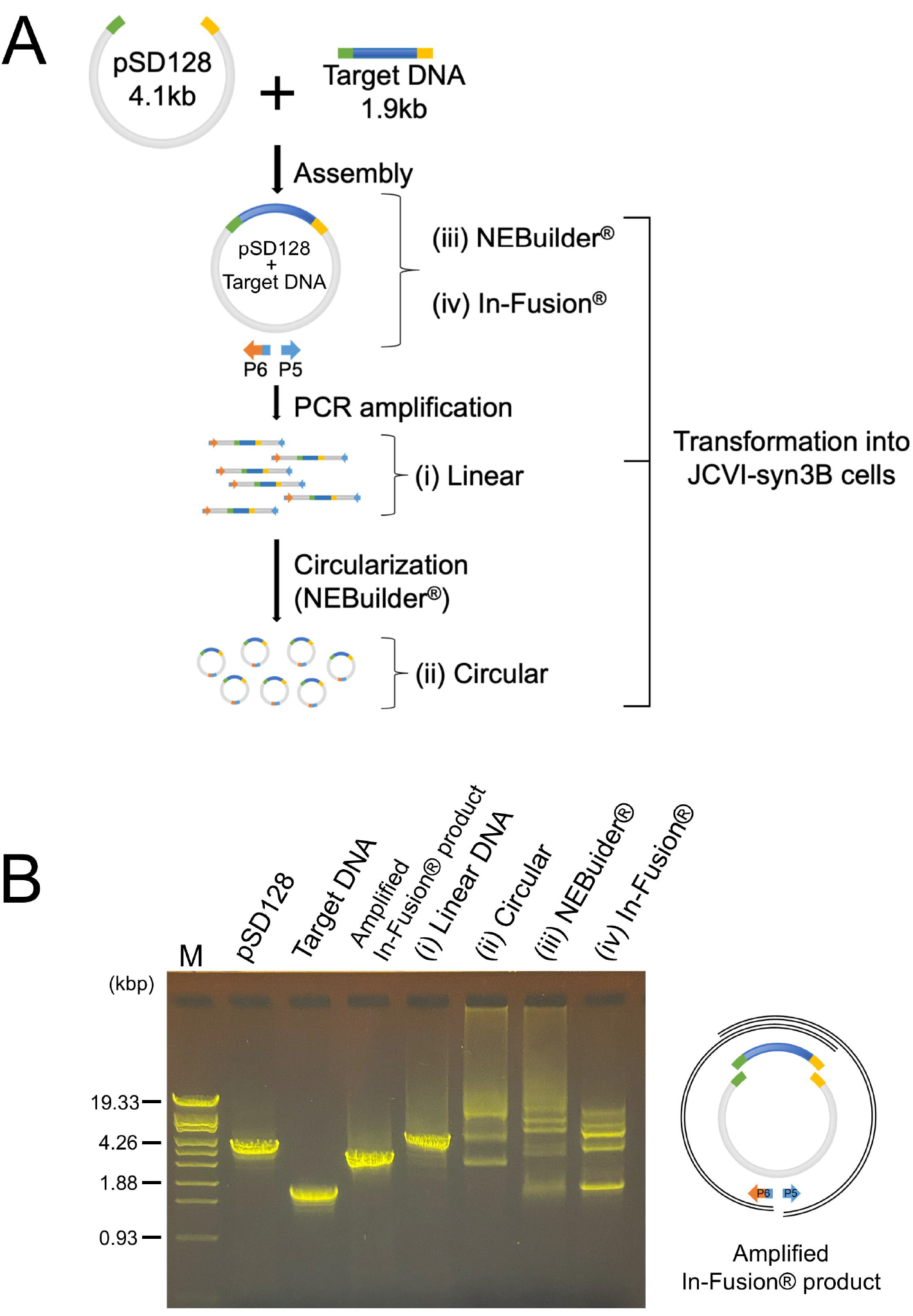
DNAs used for transformation. (A) Schematic illustration for preparation of four DNAs. (i) Linear DNA, including the target DNA, Cre/*loxP* system, and puromycin resistance gene. (ii) Circular DNA prepared using NEBuilder®. (iii) DNA assembled from two fragments using NEBuilder®. (iv) DNA assembled from two fragments using In-Fusion®. Four types of DNA were used for syn3B transformation. (B) Left: Electrophoresis of DNAs used for transformation. Size markers are shown in the left lane. pSD128 and Target DNA are the fragments to be assembled. The fourth lane is the result of PCR amplified from In-Fusion® product. Four DNAs used for transformation are shown as numbered (i) – (iv). 200 and 150 ng DNA loaded on lanes (ii) – (iii) and (iv), respectively. The other lanes were loaded with 100 ng DNA. Right: Schematic illustration of amplified In-Fusion® product. The outermost arcs present DNA fragments amplified by this PCR.

First, we tried an assembled and amplified DNA fragments (Fig. 2(i)). Initially, we performed assembly using the In-Fusion® system, but the size of the resulting target fragments was shorter than the full length (Fig. 2B). In-Fusion® system has no DNA polymerase and ligase for the gap-filling activity that prevents mutagenesis during filling [18,19]. Therefore, the target DNA with the region between P5 or P6 should be amplified by the PCR (Fig. 2B right). This assumption was supported by observations that the amplified fragment changed with the length of target fragment. Subsequently, we used NEBuilder® with an in-built gap-filling activity [20]. Using NEBuilder®, we successfully amplified the DNA fragments (Fig. 2B). Upon transforming syn3B with 100 ng of the PCR product, 392 ± 124 colonies (n = 6) were obtained. However, the efficiency was approximately 1/50 of that achieved with a plasmid (pSD128 including target DNA) isolated from *E. coli* (Fig. 3A(i)). We investigated the cause for this low efficiency by performing transformation using circularized DNA.

**Fig. 3.**
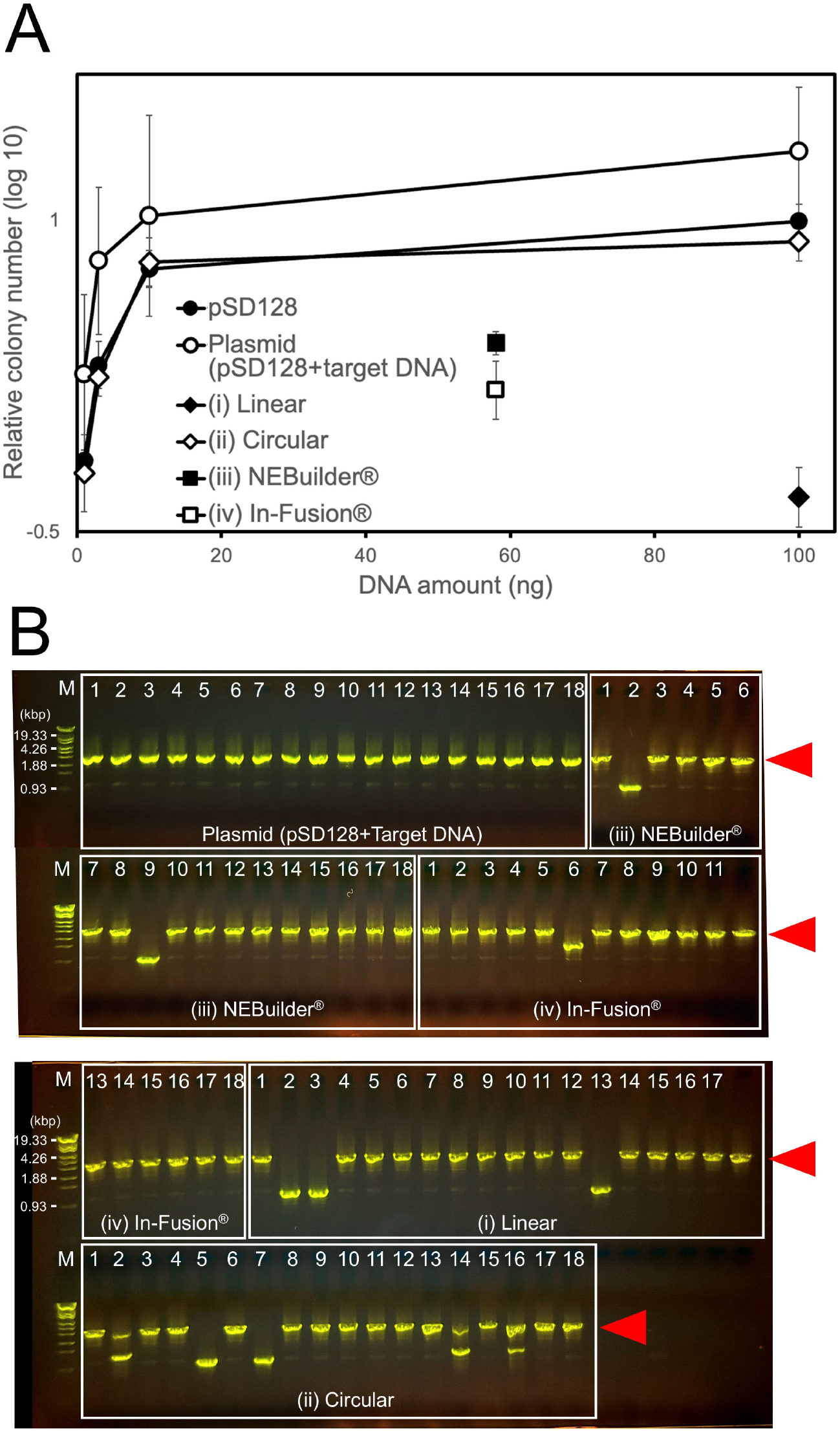
Transformation of syn3B. (A) Efficiencies with various DNAs, shown as values relative to the results of 100 ng pSD128, ranging 1,800∼23,000 colonies per mL. Amount of circular DNA was estimated based on the electrophoresis band intensity. DNA amounts obtained with (iii) NEBuilder® and (iv) In-Fusion® accounted for the total DNA amount used for assembly. (B) Direct PCR analysis of the transformants. PCR was performed using P7 and P8 primers as shown in Fig. 1A on 18 colonies obtained using individual DNAs. The band position for expected transfer is marked by red triangles.

### Transformation using assembled, amplified, and circularized DNA

PCR products were circularized by NEBuilder® (Fig. 2(ii)). The amount of DNA converted to a form other than the original linear form was estimated from the agarose gel electrophoresis band intensity (Fig. 2B), which indicated a 94.6% conversion. The DNA mixture was used for transformation (Fig. 3A(ii)). Notably, 6,458 ± 1428 colonies from 425 µL cultures (n = 6) were obtained. The transformation efficiency was approximately 3/10 that achieved with the plasmid isolated from *E. coli*. These results suggest that circularized DNA achieves a transformation efficiency similar to that achieved with the supercoiled plasmids isolated from *E. coli*. Next, we examined the products assembled without PCR amplification.

### Transformation using assembled circular DNA

The product assembled using NEBuilder®, which was expected to be circular, was directly transferred into the syn3B genome (Fig. 2(iii)). The DNA product assembled using 30 ng vector and 28 ng target DNA was transferred into syn3B, resulting in 2,067 ± 262 colonies from 425 µL cultures (n = 6; Fig. 3A(iii)). This shows the advantage of circular DNA in the uptake process. Next, we assembled the product using In-Fusion® (Fig. 2(iv)) and obtained 1,282 ± 418 colonies from 425 µL cultures (n = 6; Fig. 3A(iv)), indicating plasmid instability caused by the gap region (*P* = 0.005 via Student’s *t*-test). Here, the transformation efficiency could not be directly compared with that of the conventional method as the precise amount of assembled DNA was unknown.

However, sufficient number of colonies were obtained for gene transfer using the NEBuilder® and In-Fusion® products.

### Unexpected recombination in the transformants

We examined the transfer of the target DNA into *loxP* via direct PCR using primers P7 and P8 (Fig. 1) for 18 transformants, and repeated the whole process of this examination for six times (Fig. 3B). For the colonies obtained using the conventional method, we observed PCR bands at the correct positions in 97% of colonies. However, compared to that of the conventional method, success rates (74, 75, 93, and 92%, n = 108) were low for the linear (Fig. 2(i)), circular (Fig. 2(ii)), NEBuilder®-assembled (Fig. 2(iii)), and In-Fusion®-assembled (Fig. 2(iv)) DNAs, respectively. This decrease in success rate may be due to unexpected amplification during PCR, because the decrease depended on number of PCR application and the success rate was 97% when the plasmid cloned in *E. coli* was used. Additionally, transformation efficiency of our direct transformation method was lower than that of conventional transformation using plasmids isolated from *E. coli*.

Originally, we started this method to overcome problems in transferring AT-rich Mollicutes DNA into the syn3B genome. To date, we have constructed 58 syn3B transformants with AT content of 65–76%. Moreover, using the established method, we successfully transferred a 14.5-kb fragment consisting of eight genes of *M. mobile* into the syn3B genome. We also transferred the genes of *Haloplasma contractile* into syn3B using the new method [21,22], which is impossible with *E. coli*-based gene manipulation systems. Approximately 58 kb of the transferred DNA region was sequenced from 18 transformants. Two clones each with single-nucleotide deletions and substitutions were observed. These mutations were possibly caused by PCR, as similar mutations have been reported in *E. coli*-based recombination of PCR products.

Cre/*loxP* system is used for genome and gene manipulation widely for microorganisms, animals, plants, and human [23-25]. Although our study focused on basic research of syn3B, the direct assembly method may be applicable for other targets.

## Conclusion

Overall, we developed an *in vitro* direct rapid method for gene transfer to the JCVI-syn3B strain appreciable for Cre/*loxP* recombination. Our method is particularly useful for DNA sequences unstable or harmful to *E. coli*, and also appreciable for various systems based on Cre/*loxP* recombination.

## Author Contributions

AU performed all of the experiments. AU and MM wrote an early version of the manuscript. All authors contributed to the project and manuscript.

## Data Availability Statement

All data generated/analyzed in this study are included in this article.

## Acknowledgements

This study was supported by the Grant-in-Aid for Scientific Research (A) (grant number JP17H01544), JST CREST grant (grant number JPMJCR19S5 to MM), and Grant-in-Aid for JSPS Fellows (grant number JP24KJ0189 to HK).

